# Analysis of High Molecular Weight RNA-induced silencing complex (HMW-RISC) in CD8^+^ T cells identifies miR-7 as a modulator of T cell activation

**DOI:** 10.1101/2020.05.17.100339

**Authors:** Matilda Toivakka, Katrina Gordon, Sujai Kumar, Rose Zamoyska, Amy Buck

**Affiliations:** Institute of Immunology & Infection Research, School of Biological Sciences, University of Edinburgh, Edinburgh, EH9 3FL

## Abstract

Increasing evidence suggests mammalian Argonaute (Ago) proteins partition into distinct complexes within cells, but there is still little functional understanding of the miRNAs differentially associated with these complexes. In naive T cells, Ago2 is found almost exclusively in low molecular weight (LMW) complexes which are associated with miRNAs but not their target mRNAs. Upon T cell activation a proportion of these Ago2 complexes move into a newly formed HMW RISC which is characterized by the presence of the GW182 protein that mediates translational repression. To identify the miRNAs expected to be potent in suppressing targets, we followed HMW RISC formation upon activation of CD8^+^ T cells. We show that while most miRNAs distribute between HMW and LMW RISC in activated T cells, several miRNAs were dominant in one complex over the other. Among these, miR-7 is enriched in HMW RISC and inhibition of miR-7 upon T cell activation leads to increased production of IL-2 and expression of IL-2-regulated proteins including the α-subunit of the IL-2 receptor, CD25; transferrin receptor, CD71; and amino acid transporter, CD98, which are direct miR-7 targets. Our data support a model where recruitment of miR-7 to HMW RISC restrains IL-2 signalling and the metabolic processes regulated by IL-2 and thus modulates T cell activation.

## INTRODUCTION

miRNAs are ∼22 nucleotide small non-coding RNAs that bind to Argonaute (Ago) proteins within RNA induced silencing complexes (RISC). RISCs can form either low molecular weight (LMW) or high molecular weight (HMW) complexes depending on their interactions with other proteins (Höck et al., 2007; Landthaler et al., 2008; Olejniczak et al., 2013). HMW RISCs include mRNA binding proteins, RNA helicases, ribosomal proteins and proteins involved in gene silencing such as the trinucleotide repeat containing 6 (TNRC6) family of proteins. GW182 (TNRC6A), TNRC6B and TNRC6C are thought to be functionally redundant and bind Ago2 through the N-terminus whereas the C-terminal domains recruit other mRNA binding proteins and metabolism factors, including poly(A) binding proteins, deadenylases and decapping proteins that cause translational repression and degradation of the target mRNA (Behm-Ansmant et al., 2006; Eulalio et al., 2008; Fabian and Sonenberg, 2012; Lazzaretti et al., 2009; Lian et al., 2009). Accordingly HMW RISC was shown to be associated with ribosomes and contained the machinery required for miRNA target suppression (La Rocca et al., 2015).

In contrast to most cell lines, resting primary cells, including naive T cells, contain predominantly LMW RISC complexes and express little if any GW182 protein (La Rocca et al., 2015). Upon stimulation by mitogenic signalling (Olejniczak et al., 2013) or T cell receptor (TCR) signalling (La Rocca et al., 2015) the GW182 protein is induced and interacts with Ago protein to promote the formation of HMW RISC. On the premise that only HMW RISC complexes contain miRNAs that are engaged with mRNA targets, we sought to identify miRNAs associated with HMW RISC and study their functions in CD8^+^ T cells. Using OT-I TCR transgenic mice we stimulated CD8^+^ T cells with their physiological peptide ligands and followed the induction of GW182 protein and the formation of HMW RISC. Sequencing of HMW RISC-associated miRNAs identified miR-7 as a key miRNA in CD8^+^ T cells that functions to modulate production of Interleukin-2 (IL-2) and proteins downstream of IL-2 that are involved in T cell growth and nutrient uptake.

## RESULTS

### Ago2 forms HMW RISC with GW182 in activated CD8^+^ T cells

In order to examine miRNA expression and formation of RISC during CD8 T cell activation, we utilised OT-I CD8^+^ T cells that express an ovalbumin-reactive transgenic T-cell receptor (TCR) and respond to the ovalbumin-derived agonist peptide SIINFEKL (N4). Antigen stimulation drives activation of naive CD8^+^ T cells, promoting transcription and translation of many genes and proteins, including transient expression of the cytokine IL-2, which plays important roles in promoting proliferation and T cell differentiation. To sustain proliferation and to generate fully differentiated cytotoxic lymphocytes (CTLs), additional exogenous IL-2 cytokine was added to the cultures on day 2 (d2) and d4 (Fig1A).

**Figure 1.**
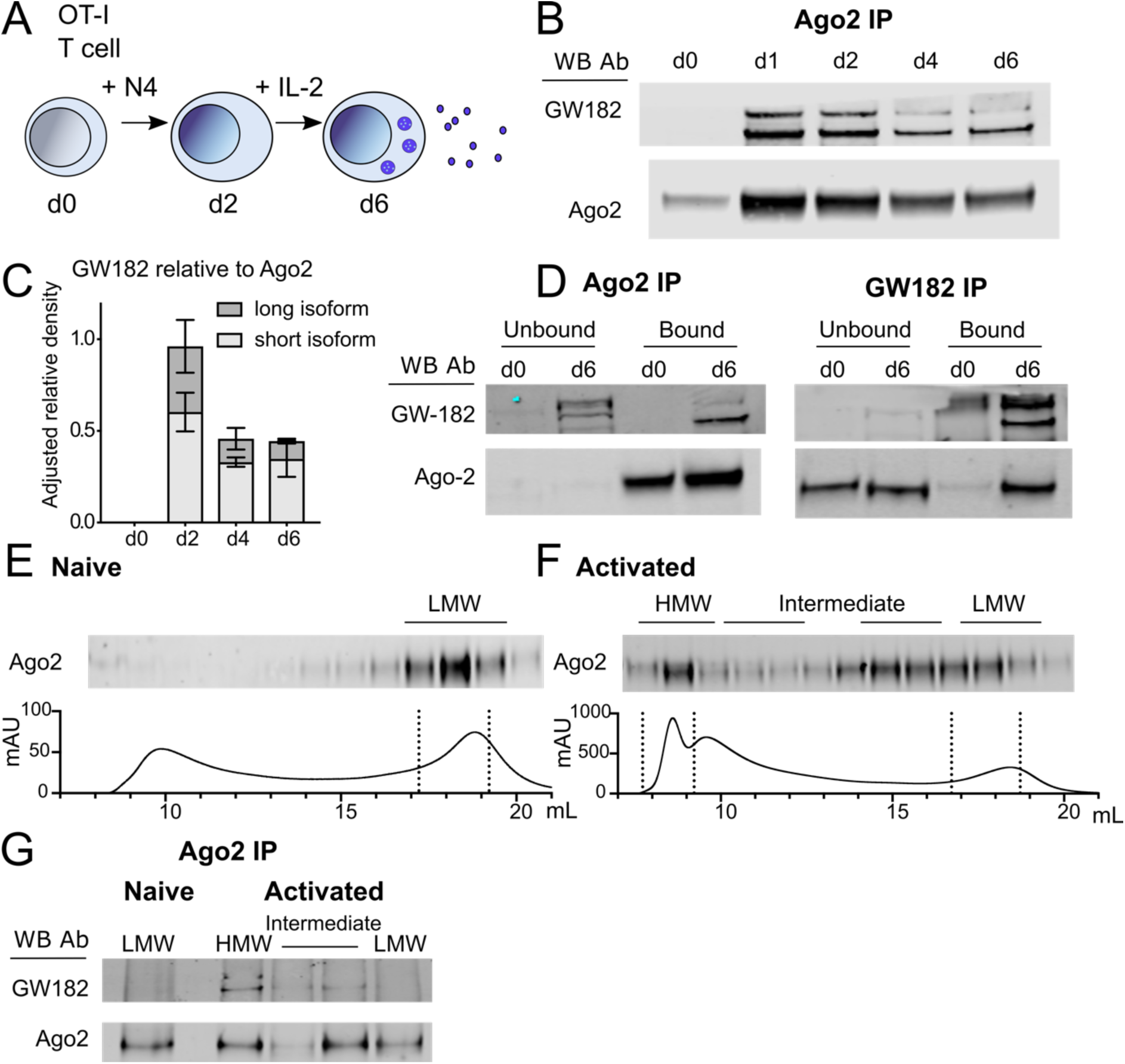
Ago2 forms HMW RISC with GW182 in activated CD8^+^ T cells. (A) Schematic of *in vitro* activation of OT-I CD8^+^ T cells with N4 and differentiation to cytotoxic effector cells with IL-2. (B-C) Ago2 was immunoprecipitated from cell lysates (2×10^7^ cells) collected from a time-course of CD8^+^ T cell activation and western blots were probed with Ago2 and GW182 antibodies (A). Quantification of the western blots measured on a Li-cor imager as GW182 band intensity relative to Ago2 band intensity, shown normalised to expression of Ago2 on d2 (B). Mean and range from 2 independent experiments. (D) Western blot from Ago2 and GW182 IPs showing bound and unbound fractions, from naive and d6 activated CD8^+^ T cell lysates (2×10^7^ cells per IP), probed with Ago2 and GW182 antibodies as indicated. (E-F) Size exclusion chromatography fractions from naive (D) and d2 activated (E) CD8^+^ T cell lysates (8×10^7^ cells). Protein elution curves show fraction volume (in mL) and absorption at 280 nm (in milli-Absorption Units). Western blot from precipitated protein from fractions probed with Ago2 antibody. (G) Ago2 IP from HMW, intermediate and LMW fractions pooled together as indicated on western blots and elution curves. Western blot of IP samples probed with Ago2 and GW182 antibodies.

To observe the dynamics of association between Ago2 and GW182, we immunoprecipitated Ago2 from naive and activated OT-I cells and detected both proteins by western blotting. GW182 co-immunoprecipitated with Ago2 from activated T cells as expected (La Rocca et al., 2015) and the two proteins remained associated during 6 days of culture, with a slight decrease in the amount of GW182 relative to Ago2 protein after day 2 (Fig1B-C). Two isoforms could be detected for GW182, with the shorter isoform predominantly immunoprecipitating with Ago2 particularly at the later time points (Fig1B-C). To further investigate this interaction, we immunoprecipitated Ago2 and GW182 from naive and activated T cells and performed a western blot of both the IP and unbound fractions. As before, Ago2 was efficiently immunoprecipitated from both naive and activated T cells, whereas only the shorter isoform of GW182 co-immunoprecipitated with Ago2 from the activated T cells (Fig1D). The reciprocal GW182 IP showed Ago2 in naive T cells remained in the GW182-unbound fraction, while activated T cells contained both GW182 bound and unbound Ago2 (Fig1D). We then used size exclusion chromatography to confirm that the GW182-bound Ago2 corresponded to a HMW RISC as described previously (La Rocca et al., 2015). As expected, all the detectable Ago2 in naive T cells was found in LMW fractions (Fig1E). In activated cells however, Ago2 was found in multiple fractions, which we define as low, intermediate and high molecular weight (Fig1F). We immunoprecipitated Ago2 from pooled fractions corresponding to HMW, two pools of intermediate fractions and LMW RISC from activated cells (as indicated in Fig1F), and LMW RISC from naive cells. As shown in Fig1G, GW182 co-immunoprecipitated with Ago2 in the HMW fractions and possibly some intermediate fractions from activated cells, but not in the LMW fractions (Fig1G). These data confirm that Ago2 and GW182 form HMW RISC in activated CD8^+^ T cells, with a fraction of Ago2 remaining GW182-unbound in LMW RISC.

### Specific miRNAs are enriched in HMW RISC

In order to determine the miRNAs associated with HMW RISC versus LMW RISC in activated CD8^+^ T cells we sequenced the small RNAs from immunoprecipitated Ago2 of each fraction (Supp. Fig1). The data were normalised to total read counts and the HMW and LMW RISC miRNAs were compared by differential expression analysis. We detected a total of 677 miRNAs of which 299 were analysed (based on criteria of average counts per million (CPM) > 4). In this dataset, 23 miRNAs were significantly enriched in HMW RISC with over 4-fold enrichment (FDR < 0.01) and 28 were enriched in LMW RISC (Fig2A, and Supplementary data 1).

**Figure 2.**
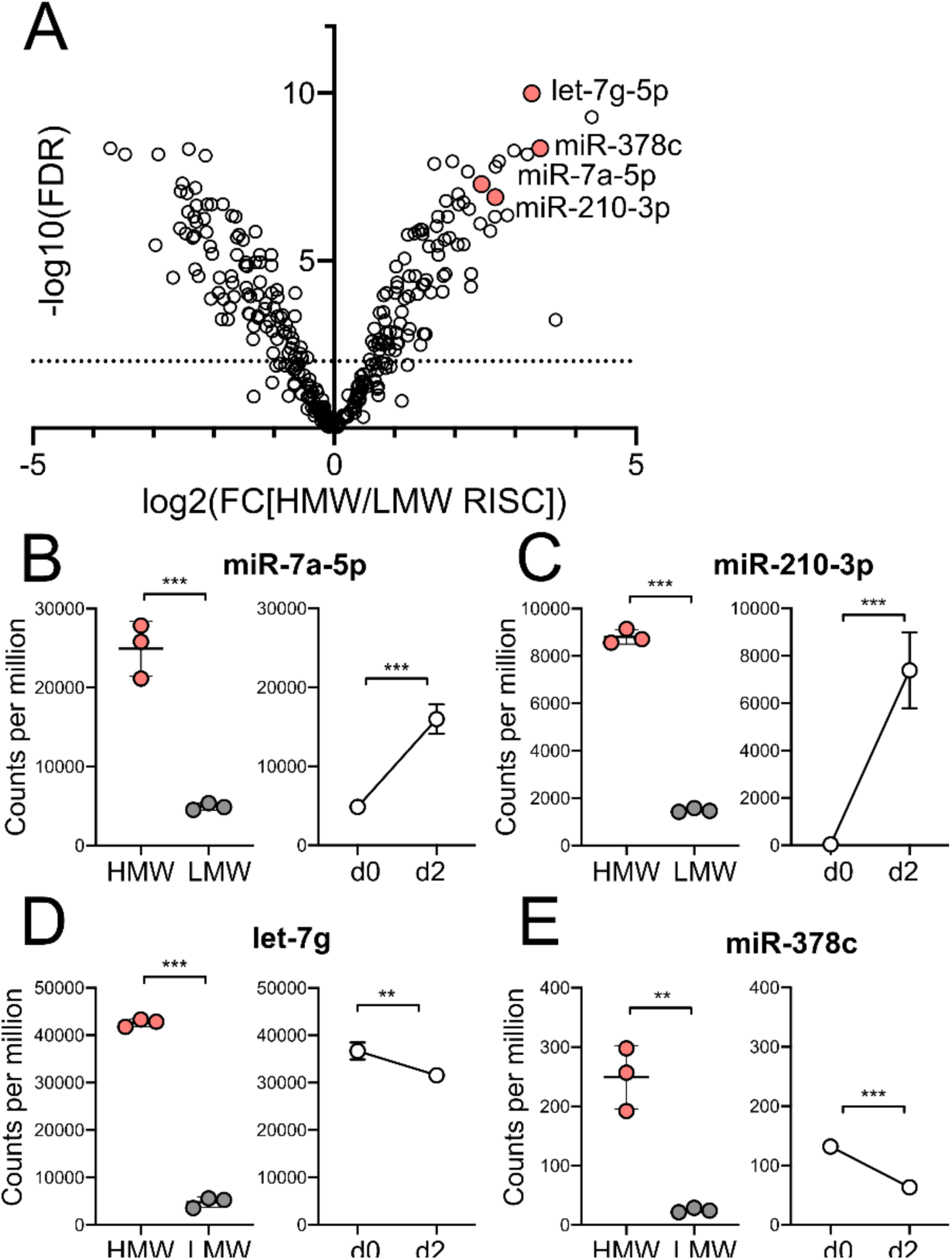
Specific miRNAs are found significantly enriched in HMW RISC in activated CD8^+^ T cells. (A) Differential expression of miRNAs between HMW and LMW RISC shown in a volcano plot of log_2_ fold change expression in HMW vs LMW RISC and -log_10_ of false discovery rate. Significant differences (FDR<0.01) above dashed line. (B-E) Counts per million (CPM) of miRNAs in HMW and LMW RISC in d2 activated cells. Expression change is shown as CPM in Ago2 IP from naive and d2 activated cells. Data are from three biological replicates in one experiment. Statistical analysis was performed using a two-tailed unpaired student’s t-test.

To compare the miRNA distributions between HMW and LMW RISC with total changes in miRNA upon T cell activation, we immunoprecipitated (IP) Ago2 from naive CD8^+^ T cells, and cells activated with peptide for 2 days (prior to addition of IL-2). As expected from published data, sequencing analysis of the IP eluants revealed dynamic changes in the miRNAs associated with Ago2 upon activation (Supp. Fig2A). We validated the upregulation of miR-155 and miR-17, and downregulation of miR-150, miR-139 and miR-181 by qPCR from total cellular RNA during a time-course of activation (Supp. Fig2B-C). These results are consistent with previous reports which showed differences in the total miRNA population in T cell subsets during activation (Rodríguez-galán et al., 2018).

At a global level the enrichments of specific miRNAs in LMW or HMW RISC did not appear to correlate with overall abundance of the miRNAs, or changes in their expression upon activation (Supp. Fig3A-C). For example, of the miRNAs enriched in HMW RISC, miR-7 and miR-210 were strongly upregulated as shown by higher CPM in the Ago2 IP from activated T cells compared to Ago2 IP from naive cells (Fig2B-C) in contrast to miR-378c and let-7g that were downregulated (Fig2D-E) upon activation. Interestingly, we observed that members of the same miRNA families characterised by a shared seed sequence displayed the same pattern in terms of enrichment in HMW versus LMW RISC (Supp. Fig3D-E).

**Figure 3.**
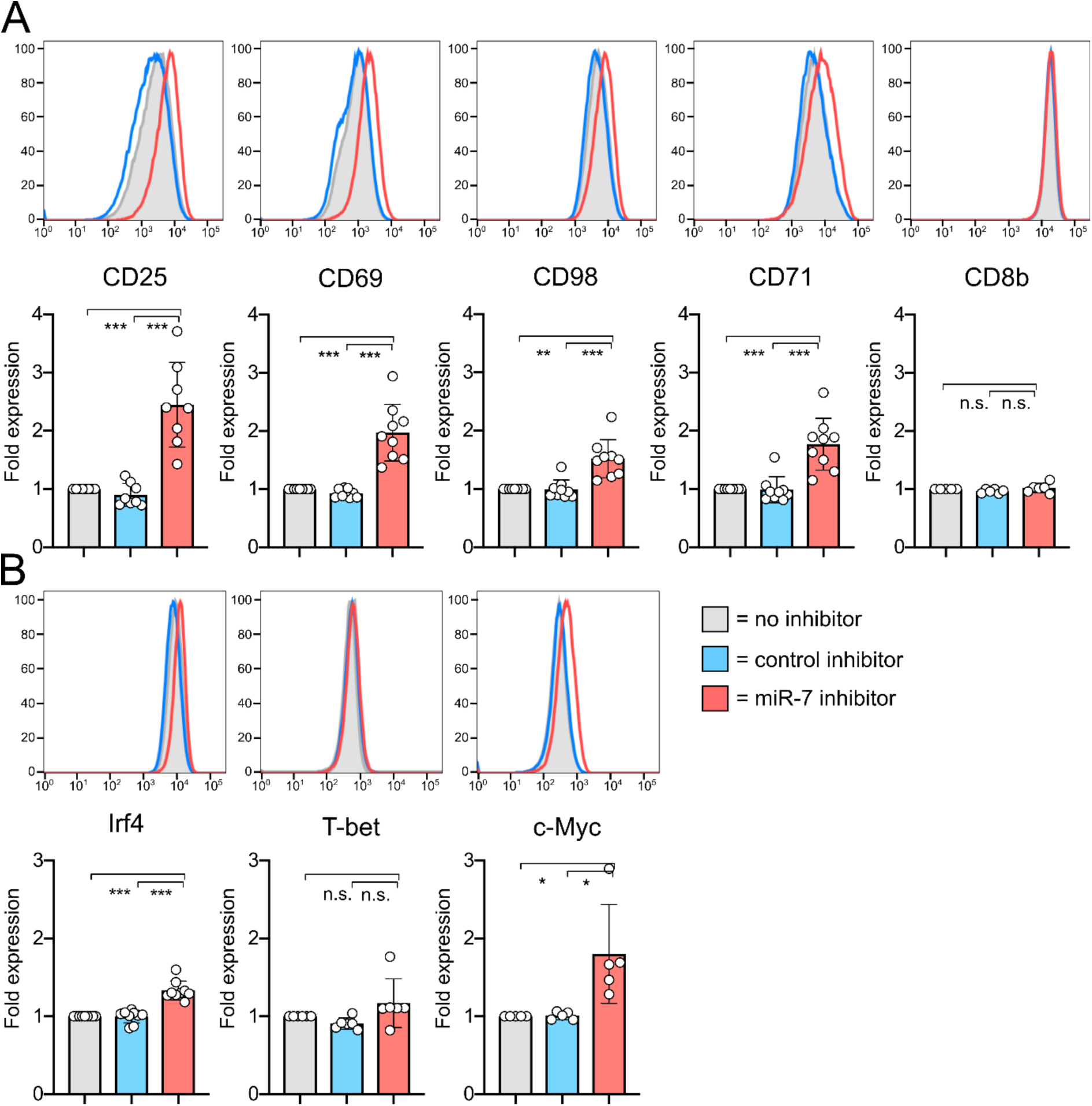
Inhibition of miR-7 during T cell activation results in altered cell phenotype. (A) Cells were activated with N4 peptide in the presence of miR-7 inhibitor and expression of surface markers (A) and transcription factors (B) was measured on d2. Representative flow cytometry histograms are shown with graphs showing individual biological replicates pooled from 4-6 independent experiments, with fold expression shown relative to ‘no inhibitor’ control. Statistical analysis was performed using a one-sample t-test to compare ‘miR-7 inhibitor’ to the hypothetical mean of 1 and a two-tailed unpaired student’s t-test to compare ‘control inhibitor’ and ‘miR-7 inhibitor’ conditions.

### Role of miR-7 in T cell activation

We hypothesised that HMW RISC enriched miRNAs would be actively engaged in target repression and focused on one of the miRNAs identified from these data, miR-7, for further investigation. miR-7 was significantly upregulated upon activation and one of the most enriched miRNAs in HMW RISC but has not been described previously to have a specific function in T cells (Gagnon and Ansel, 2019). In cancer cells however, miR-7 has been shown to be tumour-suppressive and has, for example, been shown to target components of the mammalian target of rapamycin (mTOR) pathway (Fang et al., 2011; Wang et al., 2013; Zhao et al., 2015). To study the function miR-7 in CD8^+^ T cells, we used a locked nucleic acid (LNA) inhibitor that was delivered to the naive T cells at the time of activation. Using flow cytometry, we could demonstrate efficient uptake of the fluorescently labelled miR-7 inhibitor on d1 and d2 post activation of the cells (Supp. Fig4A). Trypsin treatment prior to flow cytometry reduced the signal for the surface co-receptor CD8β but caused only a slight decrease in the fluorescent signal from the inhibitors, consistent with their internalisation (Supp. Fig4B). Cell viability was not affected by the uptake of the miR-7 inhibitor or a control inhibitor (Supp. Fig4C).

**Figure 4.**
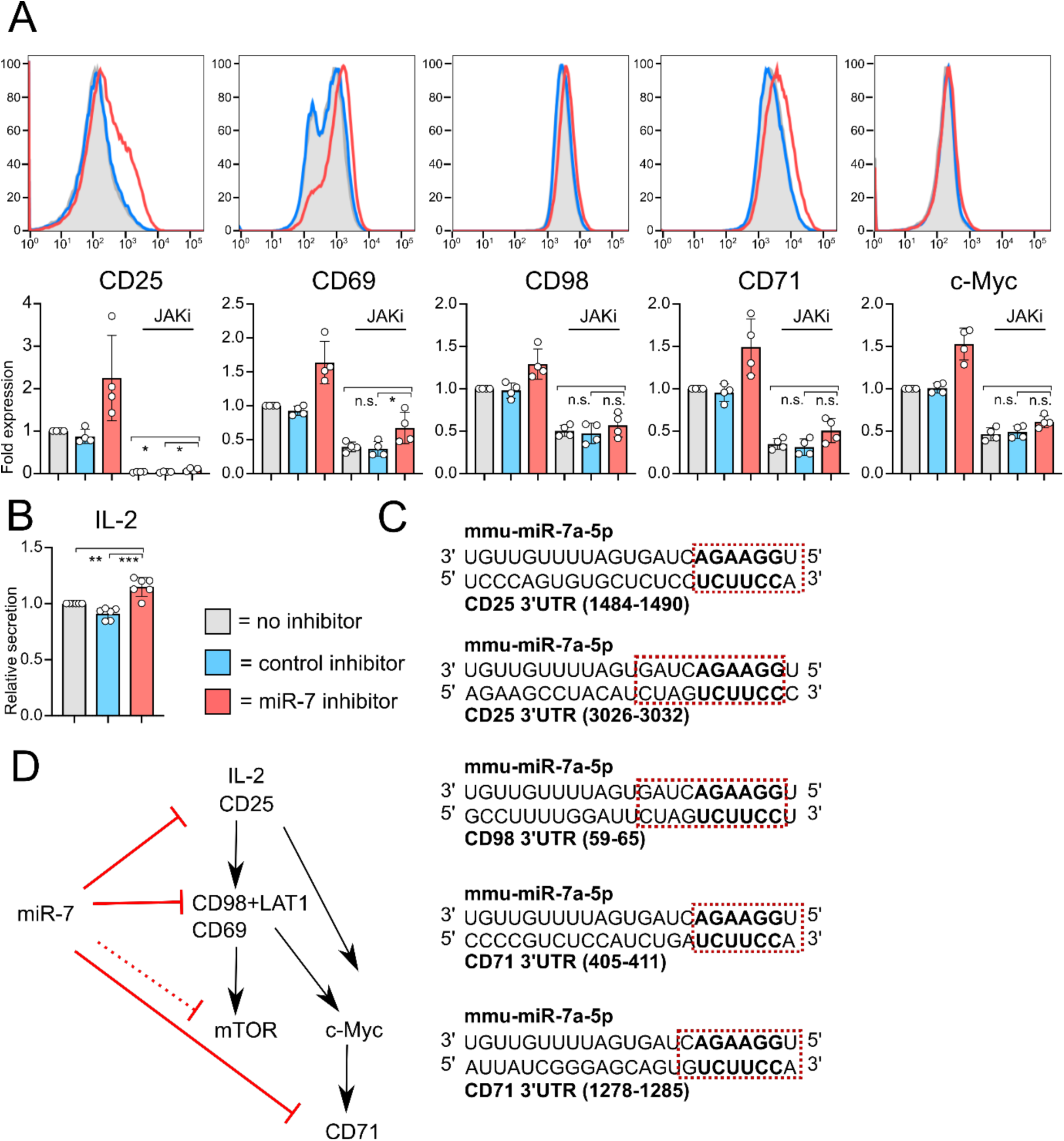
Blockade of JAK signalling only partially abrogates miR-7 inhibition. (A) Cells were activated with N4 peptide ± miR-7 or control inhibitor. JAK inhibitor (tofacitinib) was added on d1. Expression of surface markers and transcription factors was measured on d2. Representative flow cytometry histograms are shown with pooled data from 4 independent experiments. No JAK inhibitor data are reprised from Figure 4. Fold expression is shown relative to ‘no inhibitor’ control. Statistical analysis was performed using a one-way ANOVA with Tukey’s multiple comparisons test, to compare the JAK inhibitor treated conditions. (B) IL-2 secreted on d1, measured by ELISA from culture supernatants. IL-2 receptor blocking antibody was included in cultures to prevent uptake and depletion of IL-2 by the T cells. The graph is showing six biological replicates from three independent experiments, as IL-2 secretion (measured in pg/mL from culture supernatant) relative to ‘no inhibitor’ control. Statistical analysis was performed using a one-sample t-test to compare ‘miR-7 inhibitor’ to the hypothetical mean of 1 and a two-tailed unpaired student’s t-test to compare ‘control inhibitor’ and ‘miR-7 inhibitor’ conditions. (C) miR-7 target site location in 3’UTR of CD25, CD98 and CD71. The 6-mer seed site is shown in bold and other matching nucleotides are shown within the red square. (D) Schematic of proposed mechanism of action of miR-7 in CD8^+^ T cells.

We followed whether the T cell response to stimulation was altered by the inhibitors by measuring cell proliferation, cell size, expression of surface markers and of transcription factors by flow cytometry. Two days after activation, the cells had proliferated at a comparable rate (Supp. Fig4D). Naive T cells increase their size upon activation proportionally to the strength of stimulation, which can be measured by increases in forward scatter (FSC) profiles by flow cytometry. On d2 N4 stimulated cells receiving the miR-7 inhibitor were found to be slightly larger, indicative of greater cellular mass (Supp. Fig4E). We measured the expression of various receptors which are activated upon T cell stimulation and regulate cell growth, such as the IL-2 receptor alpha chain, CD25, the surface activation marker, CD69, the transferrin receptor, CD71, and the amino acid transporter, CD98, one and two days after activation in the absence or presence of the inhibitors. On d2 the cells receiving the miR-7 inhibitor specifically expressed higher levels of CD25, CD69, CD71 and CD98 whereas expression of a control surface receptor, CD8β, was unchanged (Fig3A). We also measured the expression of the transcription factors Irf4, T-bet and c-Myc, that regulate CD8^+^ T cell activation, metabolism and differentiation to effector cells. The cells receiving the miR-7 inhibitor consistently showed increased expression of c-Myc, and to a lesser extent Irf4, with no change in the expression of T-bet (Fig3B).

Naive T cell activation is a multistep process with the initial signals being delivered by engagement of the TCR and co-stimulatory molecules, while subsequent growth and effector cell differentiation is largely driven by the response to cytokines such as IL-2 (Ross and Cantrell, 2018). Since inhibition of miR-7 had no effect on the activation phenotype of the T cells on d1 (data not shown) but had strong phenotypic changes at d2, we hypothesised that miR-7 may be important in downregulating the IL-2 pathway. Although the CD25 component of the high affinity IL-2R is induced by TCR stimulation initially, its expression is subsequently maintained by, and proportional to, the availability of IL-2 to the cell (Kim et al., 2006). Both TCR signals and IL-2 contribute to the upregulation of CD98 that interacts with LAT1, and potentially CD69, to form the system L (‘leucine preferring system’) amino acid transporter responsible for large neutral amino acid (LNAA) uptake in activated T cells (Cibrian et al., 2016; Sinclair et al., 2013). Key metabolic regulators such as the mTOR complex 1 (mTORC1) and c-Myc, that regulates the expression of glucose, glutamine and transferrin receptors, are also regulated by IL-2 signalling and system L amino acid uptake (Howden et al., 2019; Preston et al., 2015; Rollings et al., 2018).

To determine the effect of miR-7 on IL-2 signaling we used Janus kinase (JAK) inhibitor tofacitinib, which blocks signalling downstream of the IL-2 receptor. The addition of the JAK inhibitor on d1 caused a complete downregulation of CD25 and a strong reduction in the expression of CD69, CD71 and CD98, as expected since IL-2 is important in maintaining production of these proteins (Supp. Fig4F) (Rollings et al., 2018). Addition of the JAK inhibitor lessened the phenotypic differences between miR-7-inhibited and control cultures (Fig4A) and was consistent with a slight but statistically significant increase in IL-2 production which was observed in the presence of the miR-7 inhibitor (Fig4B). These data indicate that miR-7 dampens the production of IL-2 following TCR stimulation, which in turn modulates the expression of a number of downstream molecules by d2 of activation. However, following inhibition of JAK; CD25, CD69, CD71 and to a lesser extent CD98 and c-Myc were still more highly expressed in the presence of the miR-7 inhibitor (Fig4A), consistent with the 3’UTRs of most of these genes containing binding sites for miR-7 (Fig. 4C) and previous validation of miR-7 targets CD71 and CD98 in cell lines (Miyazawa et al., 2018; Nguyen et al., 2010). Together these findings indicate that miR-7 regulates the pathways controlling IL-2 signalling in CD8^+^ T cells on multiple levels, impacting IL-2 production, expression of the IL2 receptor CD25, amino acid uptake through CD98 and transferrin uptake by CD71 (Fig4D).

## DISCUSSION

The abundance of a miRNA in a cell has historically been taken as an indication of its functional importance. However many recent observations across plant and animal systems do not fit this dogma and suggest miRNAs can exist in distinct RISCs with different functional properties (Dalmadi et al., 2019; Flores et al., 2014; Olejniczak et al., 2013; Powell et al., 2020; La Rocca et al., 2015; Wu et al., 2013). In T cells HMW RISC association was previously shown to correlate with increased miRNA suppressive capacity: members of the let-7 and miR-16/15 families were shown to be more repressive for a reporter construct in activated, compared to naive T cells, despite downregulated expression (La Rocca et al., 2015). Consistent with these studies, by sequencing HMW and LMW RISC associated miRNAs from activated CD8^+^ T cells we found that, while most miRNAs could be detected from both complexes, some were dominant in one over the other. This suggests that the importance of a miRNA in canonical suppression of mRNA targets may have more to do with its association with HMW RISC than its overall abundance in the cell. We demonstrate that one of the most enriched miRNAs in HMW RISC is miR-7 and show for the first time that this miRNA plays a key role in regulating T cell activation. This regulation acts on CD25, CD69, CD71, CD98, c-Myc and Irf4, which are all interlinked in their roles in sustaining T cell activation through promoting amino acid uptake and signalling downstream of the IL-2, c-Myc and mTOR pathways (Fig4C-D). We show that CD25, CD71 and CD98 contain direct seed-based target sites for miR-7 in their 3’UTRs, and speculate that additional targets may exist to explain the altered production of IL-2 itself. miR-7 therefore has the potential to restrict IL-2 production and several molecules downstream of IL-2 signalling to limit the extent of T cell activation. This could be important in conditions of persistent inflammation to avoid T cell exhaustion (Gaud et al., 2018; Wherry and Kurachi, 2015).

Interestingly, in T cells GW182 expression and HMW RISC formation is at least partly regulated by mTOR signalling following T cell activation (La Rocca et al., 2015). We propose therefore that formation of HMW RISC following mTOR signalling is one feedback pathway to regulate T cell activation through miR-7 and potentially other miRNAs. Indeed the formation of HMW RISC is likely to be important to regulate signalling in a range of other contexts and is dynamically regulated by mitogenic signals, with the deprivation of growth factors or glucose causing decreased expression of GW182 and formation of a LMW RISC in cell lines (Olejniczak et al., 2013). We do not rule out that LMW RISC could have functional properties, for example to direct cleavage of targets that are highly complementary to miRNAs in Ago2, which has endonuclease activity. However this requires further characterization in T cells.

The question of if, or how, miRNAs are selectively recruited to HMW RISC is not understood currently. In CD8^+^ T cells the presence in HMW versus LMW RISC did not depend of the total abundance of the miRNA nor correlate with the level of miRNA up– or down-regulation (Supp. Fig3A-C). The distribution of miRNAs between HMW and LMW RISC could however be influenced by the availability of miRNA targets, which changes dramatically upon T cell activation due to changes in transcription and shortening in 3’UTR of certain transcripts (Sandberg et al., 2008). Overexpression of a mRNA containing miRNA target sites has been shown to increase the RISC-association of that miRNA (Flores et al., 2014). In support of this hypothesis, we found that miRNAs with identical seed sites and overlapping targets were found together in HMW RISC (Supp. Fig3D-E). For example, members of the miR-17 family, which regulate CD8^+^ T cell proliferation and cell differentiation, were found upregulated and enriched in HMW RISC (Supp. Fig3D-E) (Khan et al., 2013; Wu et al., 2012). Similarly, miR-449a and miR-449c that share a seed site with miR-34a, were upregulated and enriched in HMW RISC (Supp. Fig3D-E). We also found that, in agreement with previous results (La Rocca et al., 2015), members of the let-7 family were downregulated overall upon T cell activation but enriched in HMW RISC (Supp. Fig3D-E). It remains possible however that other factors, including modifications of miRNAs (e.g. non-template nucleotide addition) could also influence miRNA loading to Ago and assembly of RISC (Gebert and MacRae, 2019). Ago can also be modified on various residues to regulate its subcellular localisation, association with GW182, loading and turnover of miRNA and mRNAs (Bridge et al., 2017; Golden et al., 2017; Horman et al., 2013; Rüdel et al., 2011; Zeng et al., 2008). Other RNA-binding proteins may also interact with specific miRNAs or targets to influence RISC formation (Krol et al., 2010).

In summary, we have confirmed the presence of LMW RISC in CD8^+^ T cells and the induction of HMW RISC when these are stimulated with antigenic peptide. We have shown enrichment of specific miRNAs and miRNA families in HMW RISC and demonstrated that one of these, miR-7, is a novel regulator of T cell activation and metabolism that suppresses IL-2 signalling and nutrient uptake. There remain open questions in regard to the function, composition and recruitment of miRNAs to HMW RISC, but our data show that focusing on miRNAs enriched within HMW RISC can lead to identification of new functionally relevant miRNAs, whose importance could be overlooked if only focusing on abundance levels.

## METHODS

### Mice & Primary cell cultures

All mice were maintained and bred in pathogen-free conditions at the University of Edinburgh animal facilities in accordance with the UK Home Office and local ethically approved guidelines. OT-I transgenic mice (Hogquist et al., 1994) on the Rag-1KO C57BL/6 genetic background were used, as well as OT-I mice containing a tagged Ago2 knock-in (Schmolka et al., 2018) for the experiments shown in Figures 1 and 2, and Supplementary Figures 1-3. Mice were used at 7-12 weeks of age and animal sex was not expected to have a significant influence on outcomes. Single cell suspensions were prepared mechanically from mouse lymph nodes using a 70 µm filter. Cells were grown in Iscove’s Modified Dulbecco’s Medium (IMDM, Sigma Aldrich) supplemented with 10% Heat inactivated Foetal Calf Serum (FCS, Gibco), 100 U/mL Streptomycin (Gibco), 100 µg/mL penicillin (Gibco), 2 mM L-glutamine (Gibco) and 50 µM β-mercaptoethanol. The cells were grown at 37 °C.

### T cell culture

To activate OT-I cells, 2 x 10^6^ cells/mL were grown in media supplemented with 10nM N4 for 2 days. For effector cell differentiation, cells were activated with N4 as before for two days, then resuspended at 2 x 10^5^ cells/mL in media supplemented with 20ng/mL IL-2 (PeproTech). miRNA inhibitors (miRCURY LNA miRNA power inhibitor with or without 5’-FAM, Qiagen) were added directly to culture media at the time of activation and were taken up by the cells by spontaneous translocation across the cell membrane. 500nM miRNA inhibitor (mmu-mir-7a-5p) or control inhibitor (negative control A) was added to naive T cells in cell culture media with N4, and phenotype was assessed on d1 and d2. For JAK inhibition, tofacitinib (Stratehc) was added to the cultures on d1 at a final concentration of 200nM and cell phenotype was measured on d2.

### Flow cytometry

To measure cell proliferation, cells were stained with CellTrace Violet cell proliferation kit (Thermo Scientific) prior to culturing. Flow cytometry staining was undertaken in 96-well round-bottom plates with a minimum of 200,000 cells per well. To distinguish live cells, the cells were stained with LIVE/DEAD Fixable Aqua dead cell stain kit (Thermo Scientific). For surface staining, the cells were incubated in Facs buffer (2.5% FBS and 0.05% sodium azide in PBS) with labelled antibodies, all Thermo Scientific unless otherwise indicated: anti-CD8β, anti-CD25, anti-CD69 (BioLegend), anti-CD71 (BioLegend) and anti-CD98. Intranuclear staining was done using the eBioScience Foxp3 / Transcription Factor Staining Kit (Thermo Scientific) and anti-Irf4, anti-T-bet and anti-c-Myc (Cell Signaling) antibodies. Flow cytometry was performed on MacsQuant Analyzer 10. Flow cytometry data were analysed using FlowJo 10 software. Flow cytometry data were generated within the Flow Cytometry and Cell Sorting Facility in Ashworth, King’s Buildings at the University of Edinburgh.

### ELISA

To measure IL-2 production, cells were cultured in the presence of an anti-CD25 blocking antibody (2BScientific, PC61). IL-2 was measured from culture supernatants using a mouse IL-2 ELISA kit (Thermo Scientific).

### Western blotting

Cells were lysed in lysis buffer (50mM Tris/HCl pH=7.8, 300mM NaCl, 1% Triton X100, 5mM EDTA, 10% glycerol) containing protease inhibitors (cOmplete protease inhibitor cocktail tablets, Roche), and if required, RNase inhibitors (RNasin Ribonuclease inhibitor, Promega). Cell lysates were denatured in LDS sample buffer (Thermo Scientific) containing 10% β-mercaptoethanol at 95°C. Proteins were fractionated on 4-12% Bis-Tris gels (NuPAGE, Thermo Scientific) and transferred onto PVDF membranes (Immobilon-FL, Merck) using the wet-transfer method. The membrane was incubated with an Ago2 antibody (a generous gift from Dónal O’Carrrol) or GW182 antibodies (Bethyl laboratories, A302-329A) then secondary labelled anti-mouse IgG (IRDye 800CW, Li-cor) or anti-rabbit IgG (Alexa Fluor 680, Thermo Scientific) antibodies in Odyssey blocking buffer containing 0.1% Tween-20 (Li-cor). The membrane was scanned using the Odyssey Clx imaging system (Li-cor). Western blot data were analysed using ImageStudioLite (Li-cor).

### Immunoprecipitations

Immunoprecipitations were undertaken using magnetic Protein G Dynabeads (Thermo Scientific) conjugated with anti-Ago2 or anti-GW182 antibodies. Cell lysates were incubated with the beads overnight after which the unbound fraction was collected. The beads were then washed using the following conditions: 1x LS-IP wash (50mM Tris/HCl pH= 7.5, 0.3M NaCl, 5mM MgCl2, 0.5% Triton x100, 2.5% glycerol), 2x HS-IP wash (50mM Tris/HCl pH=7.5, 0.8M NaCl, 10mM MgCl2, 0.5% Triton x100, 2.5% glycerol), 1x LS-IP wash, 1x PNK wash (50mM Tris/HCl pH=7.5, 10mM MgCl2, 0.5% Triton X100, 50mM NaCl). The IP-bound fraction was eluted by resuspending the beads in LDS sample buffer (Thermo Scientific) containing 10% β-mercaptoethanol and incubating at 70°C for 10 min with shaking.

### Size exclusion chromatography

Cells were flash-frozen in liquid nitrogen and lysed in 0.5 mL Superose 6 buffer (150mM NaCl, 10mM Tris/HCl pH=7.5, 2.5mM MgCl2, 0.01% reduced Triton x100, 1mM DTT) for 20 min. The lysate was cleared by centrifugation at top speed and filtering through a 20µm filter. Protein concentration was measured with Qubit 3.0 fluorometer (Thermo Scientific) using the Qubit Protein Assay kit (Thermo Scientific). Size exclusion chromatography was performed at the Edinburgh Protein Production Facility. The sample was loaded on the Superose 6 column, washed with Superose 6 buffer (omitting DTT) and 0.5mL fractions were collected. Protein was extracted from the fractions by trichloroacetic acid (TCA) precipitation.

### Reverse transcription and qPCR

RNA was extracted from cells in Trizol reagent (Thermo Scientific) using the Direct-zol kit (Zymo Research). The RNA was reverse transcribed to cDNA using the miScript RT kit (Qiagen). miRNAs were quantified using miScript Primer Assays (Qiagen) and QuantiText SYBR Green PCR Master Mix (Qiagen) on a LightCycler 480 Instrument II (Roche). qPCR data were analysed in Microsoft Excel. Data were first normalised to snRNA U6, then calculated as fold change to naive, using the ΔΔCt value method.

### Small RNA library preparation

For small RNA library preparation, RNA was isolated from Ago2-IP samples, as well as the IP input and unbound samples. The samples were resuspended in QIAzol lysis reagent and RNA was extracted using the RNeasy kit (Qiagen) and eluted in 100µl dH_2_O. The RNA was then ethanol precipitated by adding 2.5 x volume 100% ethanol, 30µl sodium acetate and 1µl GlycoBlue Coprecipitant (Thermo Scientific) to the samples which were then incubated overnight at -20°C. The following day, the samples were centrifuged at full speed for 30 min, then washed twice with 70% ethanol. The pellets were air-dried on ice then resuspended in 10-15µl dH_2_O and quantified with Qubit 3.0 fluorometer (Thermo Scientific) using the Qubit RNA HS Assay kit (Thermo Scientific). Small RNA libraries were prepared using the CleanTag™ Small RNA Library Preparation kit (Trilink) using manufacturer’s instructions and 21 cycles. The libraries were size selected to include 145-160 bps and the input, IP-unbound and IP-bound libraries were pooled at a 1:1:2 ratio. The samples were sequenced using NovaSeq 50bp paired end sequencing at Edinburgh Genomics.

### Sequencing data analysis

RNA sequencing data were processed to create miRNA count files. To assess the presence and counts of individual miRNAs, we used the miRNA counting tool QuickMIRSeq (Zhao et al., 2017). QuickMIRSeq does its own read trimming and length selection using cutadapt, which we set to trim the 3’ adapter TGGAATTCTCGGGTGCCAAGG, the 5’ adapter AGATCGGAAGAGCACACGTCT, and choose only reads between 18 and 28 bp in length for assigning to known miRNAs. The default QuickMIRSeq settings were used for identifying mouse miRNAs. The QuickMIRSeq output file miR.filter.Counts.csv was used for all downstream analyses. The miRNA count files were then uploaded on the Degust (version 3.1.0) (Powell, 2015, DOI: 10.5281/zenodo.3258933) web tool for visualisation of differential expression analysis. The differential analysis was performed using the Voom/Limma method and visualised on Degust.

## QUANTIFICATION AND STATISTICAL ANALYSIS

Statistical analysis was performed on GraphPad Prism. Details of statistical tests used, and number of biological and/or technical replicates can be found in the figure legends. Data are represented as mean and standard deviation. Statistical significance is defined as p-value <0.05.

## Acknowledgements

M.T was supported by a doctoral training grant supported by the Biotechnology and Biological Sciences Research Council (BBSRC) BB/J01446X/1 and B/M010996/1. Research in Zamoyska lab is supported by WT205014/Z/16/Z and in the Buck lab by WT097394/Z/11/Z and Wellcome Trust-University of Edinburgh Institutional Strategic Support Fund ISSF3. We thank Prof. Dónal O’Carrrol for the mouse Ago2 antibody. This work was supported by the Edinburgh Protein Production Facility (EPPF) and the Centre Core Grants (092076 and 203149) to the Wellcome Centre for Cell Biology at the University of Edinburgh. Sequencing was carried out by Edinburgh Genomics at the University of Edinburgh. Edinburgh Genomics is partly supported through core grants from NERC (R8/H10/56), MRC (MR/K001744/1) and BBSRC (BB/J004243/1). Flow cytometry data were generated within the Flow Cytometry and Cell Sorting Facility in Ashworth, King’s Buildings at the University of Edinburgh. The facility is supported by funding from Wellcome and the University of Edinburgh.

---

**Supplementary Figure 1.**
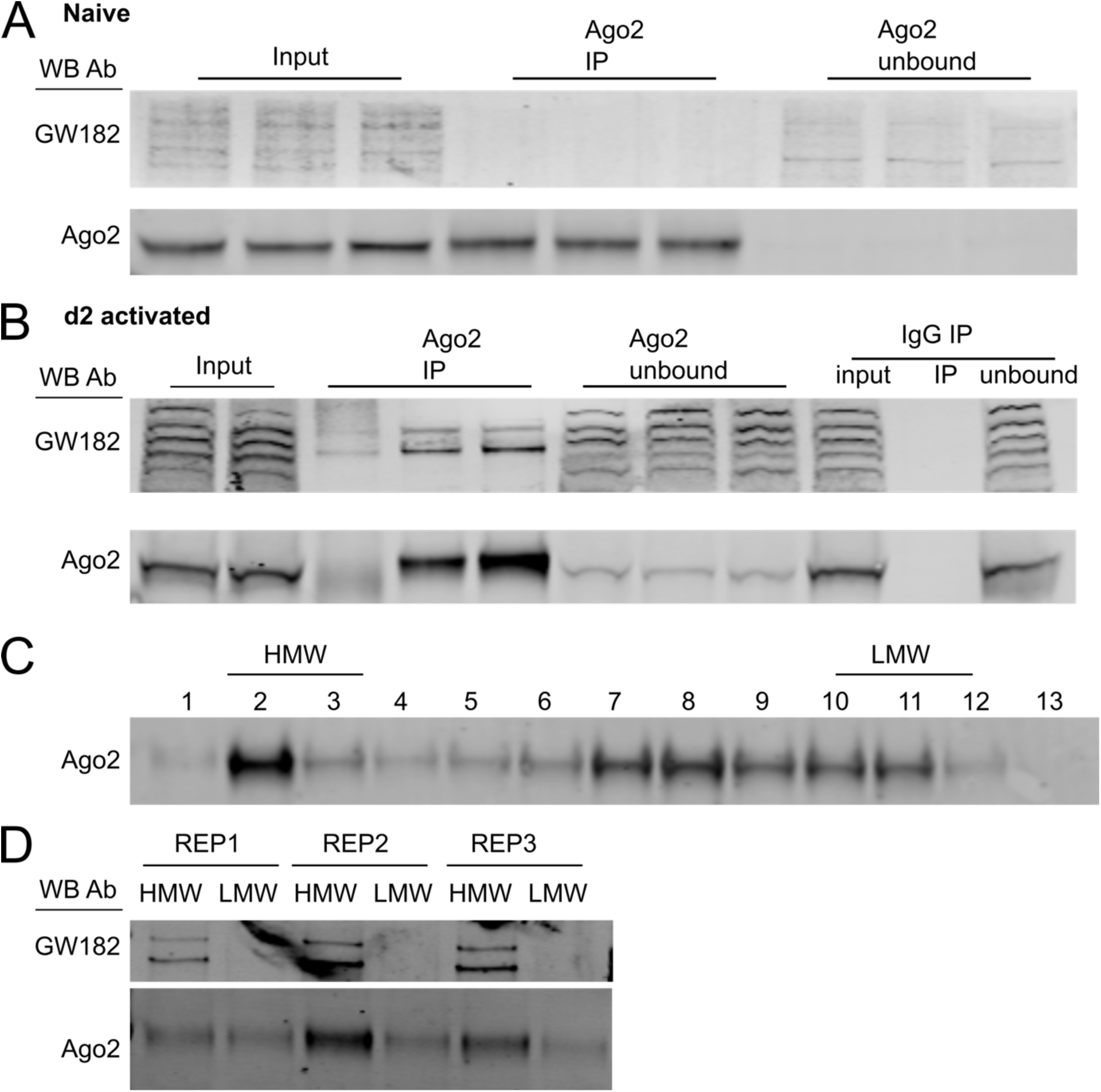
Ago2 IPs for preparation of small RNA libraries from HMW and LMW RISC. (A-B) Western blots from input, Ago2 IP (and control IgG IP) and unbound fractions from naive (A) and d2 activated (B) OT-I T cells. Blots are probed with Ago2 and GW182 antibodies. Three biological replicates are shown for naive and activated cells. (C) Western blot from protein fractions eluted by size exclusion chromatography, probed with Ago2 antibody. One replicate is shown as representative of three. (D) Western blot of Ago2 IP from pooled HMW (lanes 2-3 in C) and LMW (lanes 10-12 in C) fractions, probe with Ago2 and GW182 antibodies. Three biological replicates are shown.

**Supplementary Figure 2.**
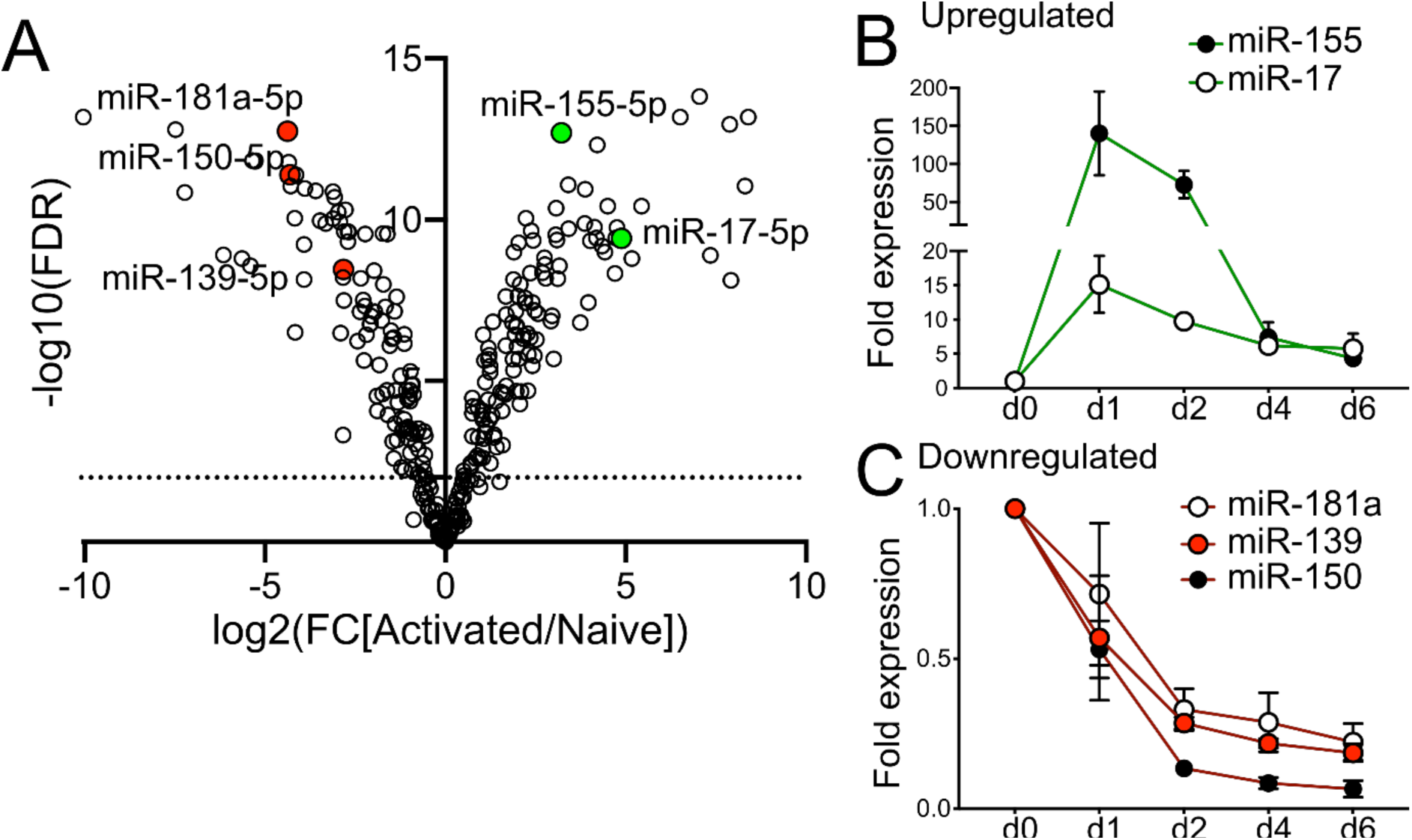
miRNA expression changes dynamically upon CD8^+^ T cell activation. (A) Differential expression of Ago2 bound miRNAs between naive and d2 activated OT-I T cells shown in a volcano plot of log_2_ fold change expression in naive vs activated cells and -log_10_ of false discovery rate. Significant differences (FDR<0.01) above dashed line. Data are from three biological replicates in one experiment. (B-C) miRNA expression measured by qPCR shown as fold expression relative to naive cells. Mean and SD from 3-4 independent experiments.

**Supplementary Figure 3.**
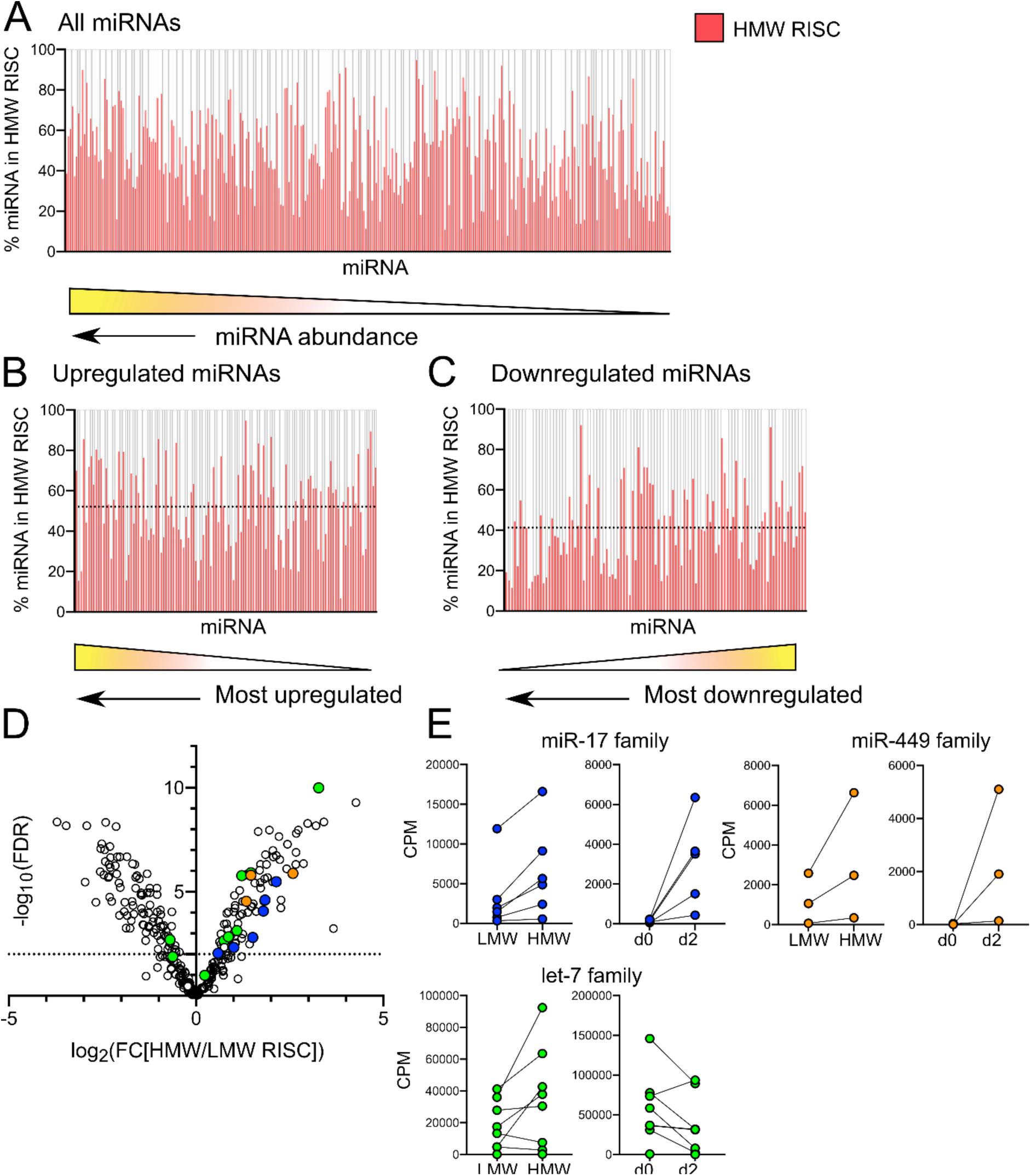
miRNA distribution between HMW and LMW RISC does not correlate with miRNA abundance. (A-C) Ratio of miRNA in HMW RISC calculated as CPM in HMW RISC over sum of CPM in HMW and LMW RISC. (A) is ranked by miRNA abundance in Ago2 IP, (B) by level of miRNA upregulation and (C) by level of miRNA downregulation. Dashed lines indicate average proportion of miRNA found in HMW RISC, 52% in (B) and 41% in (C). (D) Differential expression of miRNAs between HMW and LMW RISC shown in a volcano plot of log_2_ fold change expression of HMW vs LMW RISC and -log_10_ of false discovery rate. Significant differences (FDR<0.01) above dashed line. Members of the miR-17 family (in blue: miR-17-5p, miR-20a-5p, miR-20b-5p, miR-106a-5p, miR-106b-5p and miR-93-5p), let-7 family (in green: let-7a-5p, let-7b-5p, let-7c-5p, let-7d-5p, let-7e-5p, let-7f-5p, let-7g-5p, let-7i-5p, let-7j) and miR-449 family (in orange: miR-449a-5p, miR-449c-5p, miR-34a-5p) are highlighted. (E) CPM of miRNAs belonging to miR-17, miR-449 and let-7 families in HMW and LMW RISC in d2 activated cells. Expression change is shown as CPM in Ago2 IP from naive and d2 activated cells. Data are from three biological replicates in one experiment.

**Supplementary Figure 4.**
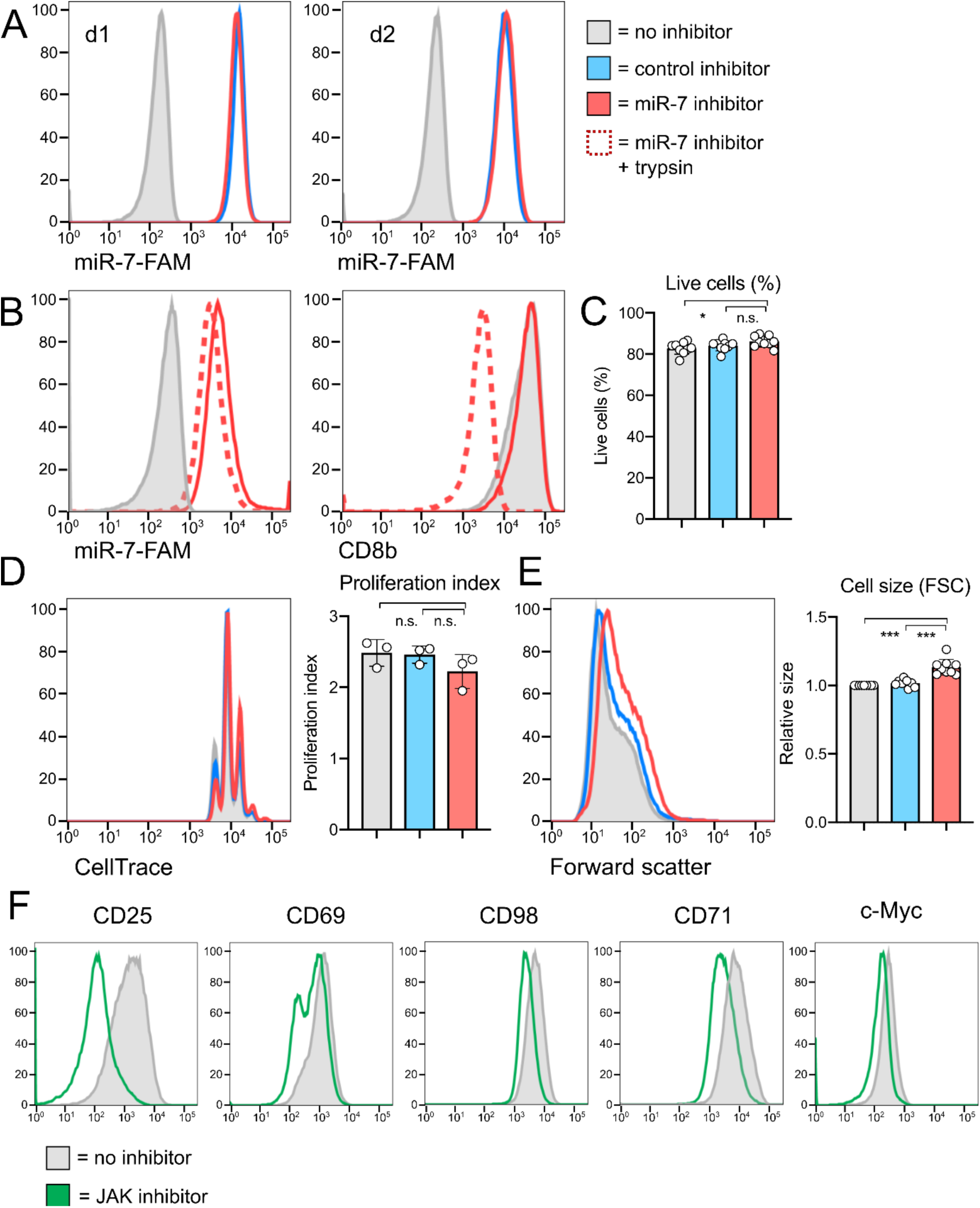
Uptake of the miR-7 inhibitor does not affect cell viability, proliferation and size. (A-B) Cells were activated with N4 in the presence of a fluorescently tagged miR-7 inhibitor. Uptake was measured by flow cytometry on d1 and d2 (A), with or without a pre-incubation with trypsin (B). (C) Percentage of live cells on d2 measured using a live/dead cell stain. (D) Cell proliferation measured by dilution of a CellTrace tracker dye on d2. Representative flow cytometry histogram is shown alongside proliferation index measured from three independent experiments. (E) Cell size measured by forward scatter in flow cytometry, shown relative to ‘no inhibitor’ control. Graph is showing individual biological replicates from 4-6 independent experiments, with fold expression shown relative to ‘no inhibitor’ control. Statistical analysis was performed using a one-sample t-test to compare ‘miR-7 inhibitor’ to the hypothetical mean of 1 and a two-tailed unpaired student’s t-test to compare ‘control inhibitor’ and ‘miR-7 inhibitor’ conditions. (F) Cells were activated with N4 on d0 and JAK inhibitor was added on d1. Expression of surface markers and transcription factors was measured on d2. Representative flow cytometry histograms (from four independent experiments) show effect of JAK inhibitor compared to no inhibitor control.

